# Feature-based valuation and its impairment in people with compulsive and addictive symptoms

**DOI:** 10.64898/2026.08.27.747558

**Authors:** Morgan G. Moll, Ryan K. Read, Thomas W. Elston

## Abstract

Effective decision-making requires evaluating options based on relevant features while reducing attention to those that are not. However, how attention prioritizes one source of information over another during value-based decisions remains unknown. Across 7 experiments, we show that irrelevant features both constructively and destructively interfered with how values are assigned to choice options. When relevant and irrelevant features agreed, choice accuracy and speed were facilitated; when they conflicted, accuracy decreased and choices slowed in proportion to the value of the irrelevant feature. We modeled this interference as a weighted sum of relevant and irrelevant feature values. This model explained individual differences in choice behavior and, critically, tracked psychiatric symptom severity related to substance abuse and obsessive-compulsive disorder. This suggests that these symptoms may, in part, reflect an impaired ability to weigh relevant against irrelevant information when assigning value to choice options – a process we term feature-based valuation.

---

Attention fundamentally shapes the choices people make. Because the brain can only process a small fraction of the sensory information available at any moment ^1–3^, it must identify and prioritize the sensory features that predict reward while suppressing attention to those that do not. Indeed, choice options that receive more attention are more likely to be chosen ^4–6^. Thus, understanding why people choose one option over another requires understanding the computations governing how the brain selects which sensory features to base choices on.

Attention is guided by multiple mechanisms ^7,8^: top-down control, bottom-up capture, and selection history. Top-down control refers to attention directed by one’s goals or intentions ^9–11^, such as looking for yellow cars when searching for a taxi. Bottom-up control refers to attentional selection that is based purely on the physical properties of a stimulus, such as brightness or contrast ^1,12,13^. Selection history refers to how stimuli previously associated with reward or punishment come to automatically and involuntarily capture attention ^14–17^. These mechanisms are thought to compete for control over attention^8^. In particular, the failure of top-down control to override selection history effects is thought to underlie how drug-related cues automatically capture attention in people with addiction ^18–23^ and, similarly, why people with obsessive-compulsive disorder (OCD) struggle to ignore environmental features that trigger their compulsions ^24–27^. How this competition is resolved therefore shapes the choices people make.

Decisions are thought to occur through a process of identifying choice options, assigning each a value, and selecting the option with the greatest value ^28,29^. Most decisions involve comparisons of options that have multiple features. When car shopping, for instance, options vary in color, size, speed, fuel efficiency, and safety rating. Which attributes matter most depends on context: a person in a rural community might prioritize a large truck with little regard for fuel efficiency, whereas a person in a city might prioritize a smaller, more efficient vehicle. Optimal choice therefore requires using attention to prioritize relevant features when assigning value to the options. Yet, attention does not always comport with our goals, especially when the context changes. For instance, people with addiction struggle to suppress attention to substance-related stimuli despite their conscious desire to ^21,30–33^. This suggests that previously reward-associated but now-irrelevant features may interfere with how the brain assigns value to choice options and, in turn, the decisions that follow. However, the computations governing how attention prioritizes one feature over another when assigning value to choice options remain unknown.

Answering this question requires bridging two traditionally siloed research areas: feature-based attention, which examines how the brain prioritizes sensory information ^34,35^, and value-based decision-making, which examines how it assigns value to choice options ^36,37^. Yet neither field has examined how attention selects which sensory features should guide value computation. If attention suppresses irrelevant features, their value should not influence choice. Alternatively, if relevant and irrelevant features are integrated during valuation, irrelevant features should intrude on choice – facilitating decisions when they align with the relevant feature and impairing choice when they conflict.

We found that irrelevant sensory information both constructively and destructively interferes with value-based choice. We show that these effects cannot be attributed to memory or task-switching effects and, rather, reflect inefficiencies in how people weigh the contribution of each feature to an option’s value. We developed a generative model that captured this interference in a single parameter – the weight given to the irrelevant feature – which provided an excellent account of individual differences in choice behavior. Finally, in a computational psychiatry study, we show that individual differences in this parameter predicted the severity of psychiatric symptoms related to OCD and substance abuse, suggesting that feature-based valuation may be a clinically meaningful dimension of cognitive control.

## Results

### Experiment 1 – measuring effect of attention during value-based choice

We recruited 50 (see **Table 1** for demographic information) human subjects to perform a feature-based valuation task designed to measure how people prioritize one sensory over another during value-based decisions. On each trial, subjects were cued to base a subsequent decision on either the color or the shape of two simultaneously presented options (**Fig. 1a**). The colors and shapes that defined each option were associated with different reward amounts (points). This created three trial types (**Fig. 1b**): *congruent*, where the best color and best shape belonged to the same option (and thus both features led to the same choice); *incongruent*, where the best color and shape belonged to different options (meaning that the best choice depended on which sensory feature was relevant); and *1-dimensional (1D)*, where only the relevant stimulus feature differed and the irrelevant feature was the same across the options (e.g. a shape trial where the choice was between two different blue shapes). This design enabled us to systematically vary the extent to which the different sensory feature-values interfered with each other.

**Figure 1.**
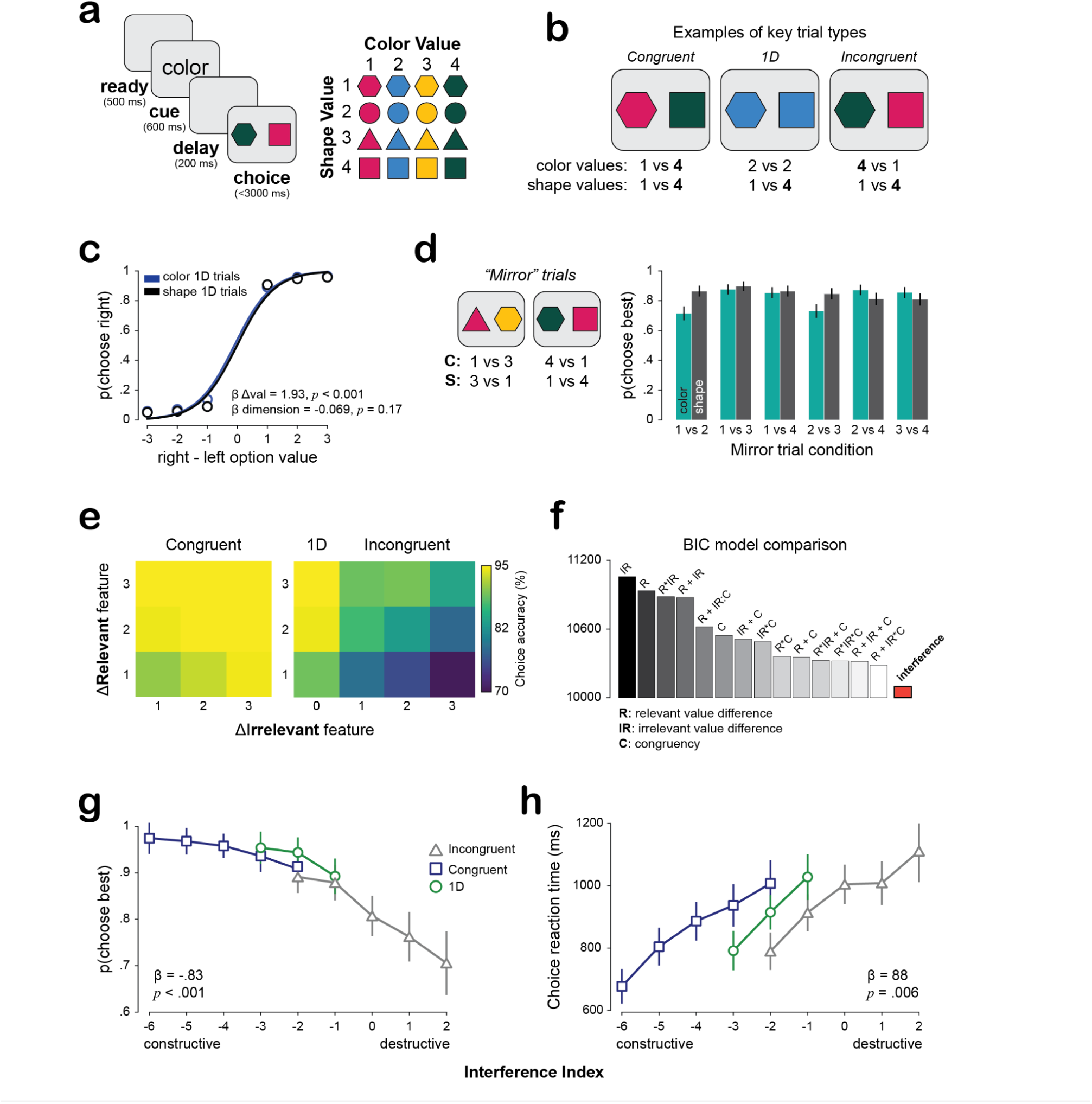
Feature-based valuation and behavioral interference effects. **(a)** Structure of the feature-based valuation task. Following an initial “ready” screen, subjects were shown a cue which indicated whether to base this trial’s decision on color or shape feature-values. After a brief delay, subjects were shown a pair of colored-shapes; the payoff of the options depended on the cue shown earlier. Following the choice response (pressing the “f” or “j” key on a keyboard), subjects were shown how many points their choice yielded. **(b)** Examples of key trial types. Congruent trials are those where the best color and shape belong to the same option.1D trials are those where the irrelevant feature is the same across both options and only the relevant feature dimension differs. Incongruent trials are those where the best color and shape belong to different options (and thus optimal choice depends on attention). **(c)** Choice accuracy during 1D trials was not different across attentional dimensions. **(d)** Examples of “mirror” trials, where the feature-values were exactly opposite. Right: choice accuracy during all mirror trial conditions. **(e)** Choice accuracy as a function of relevant and irrelevant feature-value differences during congruent, 1D, and incongruent trials. **(f)** Comparing 15 models to explain the behavior in (e) using Bayesian Information Criterion. The best-fitting and most parsimonious model was one which had a single term for the cross-schema feature-value conflict. **(g)** Choice accuracy decreased as a function of interference. The inset beta weight comes from a mixed effects logistic regression. **(h)** Choice reaction times increased as a function of interference. The inset beta weight comes from a mixed effects linear regression. Markers and error bars denote means and bootstrapped 95% confidence intervals, respectively.

**Table 1.** Participant demographics across experiments.

|  | Exp 1 | Exp 2 | Exp 3 | Exp 4 | Exp 5 | Exp 6 | Exp 7 |
| --- | --- | --- | --- | --- | --- | --- | --- |
| N (female) | 50 (21) | 50 (28) | 50 (17) | 50 (25) | 50 (23) | 50 (27) | 106 (78) |
| Mean age ± STD | 42.2 ± 14.9 | 46.3 ± 12.8 | 40.8 ± 11.4 | 43.58 ± 10.6 | 44.6 ± 11.5 | 46.1 ± 11.3 | 40.4 ± 11.7 |

Our first step was to determine whether choice accuracy differed depending on whether subjects used color or shape as the basis for valuation. For this, we focused on the *1D* trials where only the relevant feature was diagnostic for making the decision, enabling us to assess choice accuracy in the absence of any potential interference from the other stimulus dimension. As shown in **Fig. 1c**, we found a significant main effect of value difference (β = 1.93, *p* < .001, mixed effects logistic regression) but no main effect of attentional dimension (β = -0.069, *p* = .17, mixed effects logistic regression). This indicates that choice performance was comparable when subjects based their choices on color and shape.

To confirm that participants flexibly attended to the relevant (cued) feature when assigning values to choice options, we assessed choice accuracy during a subset of incongruent trials we call “mirror ” trials (**Fig. 1d**). In these trials, the relevant and irrelevant feature-value differences were exactly equal in magnitude but favored opposite options — for example, a color trial where the color values were 4 vs 1 and the shape values were 1 vs 4. If participants were simply averaging across features or ignoring the cue, performance on these trials should be at chance. Instead, participants reliably chose the option favored by the cued feature across all mirror trial conditions (all *p* < .001, paired t-tests against chance; **Fig. 1d**), confirming that they flexibly shifted their feature-based attention in accordance with the cue.

Having established that participants used the cue to guide their choices, we next examined how the agreement between relevant and irrelevant feature-values affected choice accuracy. As shown in **Fig. 1e**, participants were more likely to choose the high-value option during *congruent* versus *incongruent* trials (*p* < .001, paired *t*-test). During congruent trials, choice accuracy increased as the value-difference along the relevant and irrelevant dimensions increased. In contrast, accuracy during *1D* and *incongruent* trials systematically decreased as the value difference along the irrelevant dimension increased (note the diagonal in the figure). It is noteworthy that these effects scale with value, which suggests that irrelevant sensory information is not “filtered out” prior to valuation and influences the final choice.

To systematically determine how sensory-value interference affected choice accuracy, we compared a series of mixed effects logistic regression models which varied in how relevant and irrelevant sensory feature-values were combined (**Fig. 1f**). Prior to model comparison, we confirmed that each candidate model did not suffer from multicollinearity by assessing the variance inflation factor (VIF) for each term. We only included models where VIFs were approximately 1, indicating that the predictors were largely orthogonal to one another. We formally compared the remaining 14 candidate models using Bayesian information criterion (BIC). The best fitting model was one where the irrelevant feature values interacted with congruency – thus enabling the irrelevant information to constructively interfere with the decision variable during congruent choices and destructively interfere during incongruent choices.

Inspired by this model, we then developed an “interference index” which quantified the extent to which the relevant and irrelevant sensory features collectively “pulled” behavior towards the high value, relevant-feature option on each trial (see **Methods**). We then compared the interference index against the other 14 models and found that this single term model provided the most parsimonious account of the behavioral data (**Fig. 1f**, orange). We visualized this best model by plotting both choice accuracy (**Fig. 1g**) and reaction time (**Fig. 1h**) as a function of interference present between each trial’s sensory-value mappings. Choice accuracy decreased (β = -0.83, *p* < .001, mixed effects logistic regression) and reaction times increased (β = 88, *p* = .006, mixed effects linear regression) as interference increased. This indicates that irrelevant sensory information can both constructively and destructively influence value-based decisions.

### Interference patterns cannot be explained as an artifact of task switching

It is important to note that we did not use a block structure – color and shape trials were randomly interleaved with one another. This raises the question of the extent to which our results could be explained as an artefact of task switching. We tested this by separately analyzing and comparing switch (e.g. color→shape) and repeat (e.g. color→color) trials. While we did observe moderate performance degradation during switch as compared to repeat trial, interference effects remained significant and were above and beyond those of task switching in magnitude (**Fig. 2**).

**Figure 2.**
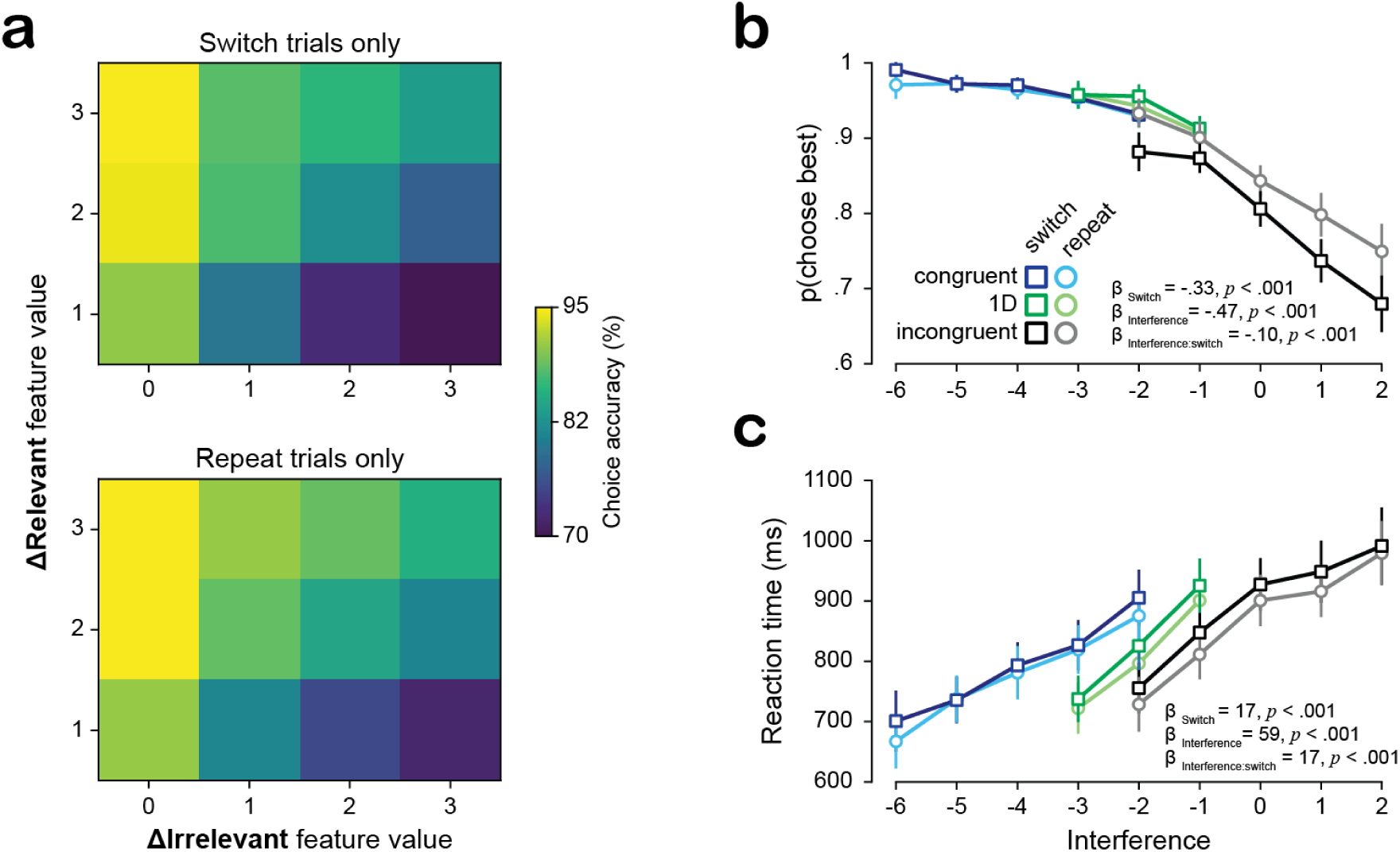
Interference effects during switch and repeat trials. (a) Choice accuracy as a function of relevant and irrelevant feature-value differences. (b) Choice accuracy as a function of cross-attentional interference, whether the trial required feature-based valuation on the same or different attentional dimension as the prior trial, and their interaction. (c) Choice reaction time as a function of cross-attentional interference, whether participants switched or repeated the same attentional dimension as the prior trial, and their interaction. Markers and error bars denote means and bootstrapped 95% confidence intervals, respectively.

### Experiments 2-4 – the roles of semantic and working memory

Another alternative possibility is that our results could reflect failures in semantic and/or working memory processing. Memorizing the discrete and arbitrary associations between color and shape in our first experiment could be challenging and so to reduce the semantic load, we made the sensory feature-value mappings continuous. Color values were organized along a gradient from blue to purple and shape values were indicated by their spikiness (**Fig. 3a**). We manipulated the working memory demand by continuously presenting the attentional cue throughout the delay and choice epochs. Thus, across 4 experiments (N = 50, each, N = 200 total), we examined how semantic and working memory individually and conjunctively contribute to interference effects during feature-based valuation.

**Figure 3.**
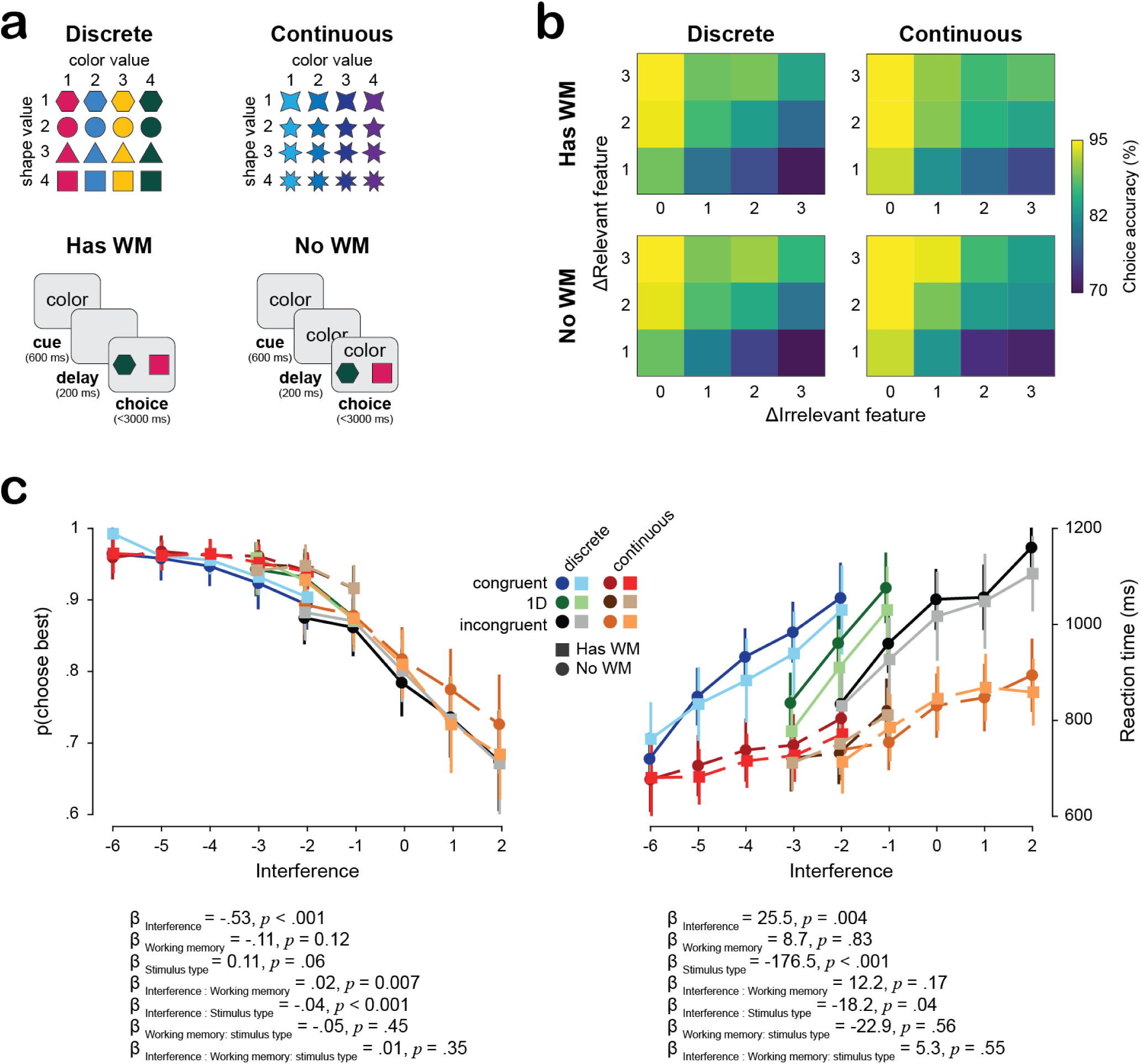
Effects of semantic and working memory on feature-based valuation. **(a)** Four experiments which varied stimulus type and working memory demand. **(b)** Incongruent and 1D choice accuracy as a function of relevant and irrelevant feature-values. **(c)** Choice accuracy and reaction time as a function of interference, stimulus type, and working memory. Markers and error bars denote means and bootstrapped 95% confidence intervals. The inset statistics denote the results of a mixed effects logistic regression for the choice data and a mixed effects linear regression for the reaction time data.

As shown in **Figure 3b**, the effect of these memory manipulations on choice accuracy during incongruent trials was minimal. We then used mixed effects regressions to measure how choice accuracy and reaction time varied as a function of interference, stimulus type, and working memory demand. We observed very strong effects of stimulus type on reaction times: participants responded much faster and were slightly more robust to interference when the stimulus features were continuous as compared to discrete. This indicates that our manipulation did indeed reduce the semantic processing burden. Across both choice accuracy and reaction time, working memory effects were marginal and often not significant. Critically, the interference main effect remained a significant, strong effect above and beyond the contributions of stimulus type and working memory (**Fig. 3c**). This indicates that the interference susceptibility we observe is likely not due to failures in semantic or working memory processing, but rather reflects a bottleneck in the ability to appropriately weigh relevant versus irrelevant sensory information when assigning value to choice options.

### Generative model of feature-based valuation

Our behavioral results demonstrate that irrelevant feature values systematically influence choice, but individuals likely differ both in how precisely they represent feature values and in how much weight they assign to relevant versus irrelevant information. To capture these sources of individual variation, we built a generative model of the decision process. The model closely parallels our interference index — which summarizes the joint contribution of relevant and irrelevant feature-value differences on each trial — but extends it in two important ways: it allows for noise in the internal representation of feature values, and it allows the relative weighting of relevant and irrelevant features to vary across individuals. Specifically, the model conceptualizes the decision variable as a weighted sum of noisy feature-values (**Fig. 4a**) and has two tunable parameters: a feature weight (ε), reflecting the relative weighting of the cued versus uncued feature, and value noise (σ), capturing stochasticity in the internal representation of reward values. The optimal strategy is to set ε = 1, allocating no weight to the irrelevant dimension. The irrelevant feature weight (1-ε) therefore directly quantifies an individual’s susceptibility to interference from irrelevant information.

**Figure 4.**
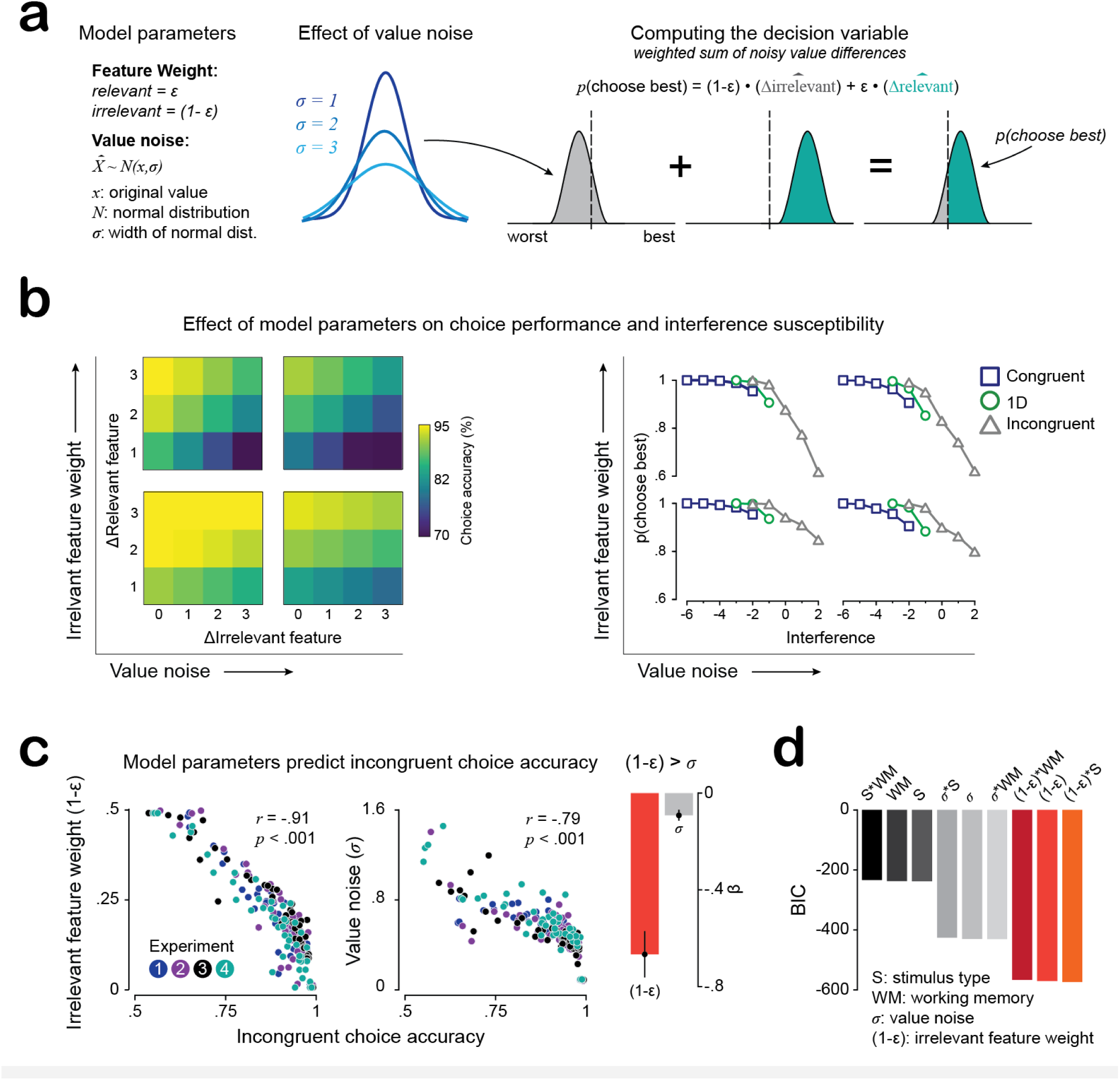
A generative model of feature-based valuation. **(a)** Schematic of model parameters and how it makes decisions. **(b)** Simulated model behavior showing the effects of modifying value noise and irrelevant feature weight. **(c)** Both value noise and irrelevant feature weight are highly correlated with individual differences in choice accuracy during incongruent trials (left). Irrelevant feature weight explains more variance in choice behavior than value noise (right). **(d)** Model comparison of factors related to irrelevant feature weight, value noise, working memory, and semantic memory on choice accuracy during incongruent trials. Models with a term for irrelevant feature weight fit best.

Our first step was to confirm whether this model was capable of producing qualitatively similar behavior to that we observe in people. We found that modifying the irrelevant feature weight in particular produced behavioral profiles very similar to what we observed in people (**Fig. 4b**). We then fit the model to each of the 200 participants in our first 4 experiments. Both model parameters were negatively correlated with choice accuracy during incongruent trials (**Fig. 4c**; irrelevant feature weight: *r =* -.91, *p* < .001; value noise (*r* = -.79, *p* < .001). We then used regression to compare which of these terms explained more variance in choice behavior. This revealed that the irrelevant feature weight was a much stronger predictor than value noise, consistent with the predictions of our earlier simulations. We then conducted a formal model comparison to assess how our memory manipulations may have interacted with our model parameters (**Fig. 4d**). The best fitting models all included a term for irrelevant feature weight. This model now provides us with a single, fittable term which captures individual differences in susceptibility to irrelevant information during decision-making.

### Experiments 5-6 – testing predictions of the model

Our modelling results indicate that individual differences in irrelevant feature weight can explain people’s susceptibility to irrelevant information during decision-making. This predicts that experimental manipulations which raise or lower irrelevant feature weight should produce correspondingly greater or reduced interference. We tested this in two experiments designed to move irrelevant feature weight in opposite directions, thereby decreasing or increasing the extent to which irrelevant information interferes with choice. To facilitate comparison to our original experiment, we used the same stimuli and timings as in Experiment 1.

In Experiment 5 (N = 50), we aimed to *decrease* irrelevant feature weight by presenting color and shape trials in blocks of 15 rather than interleaving them (**Fig. 5a**). We reasoned that this more predictable environment would let participants anticipate the upcoming relevant feature: once they inferred they were in a run of same-type trials, they could form stronger expectations about future task states and pre-commit attentional weight to the relevant dimension before each stimulus appeared. Because this anticipatory commitment draws weight away from the irrelevant feature, we predicted that blocking would reduce irrelevant feature weight and attenuate interference.

**Figure 5.**
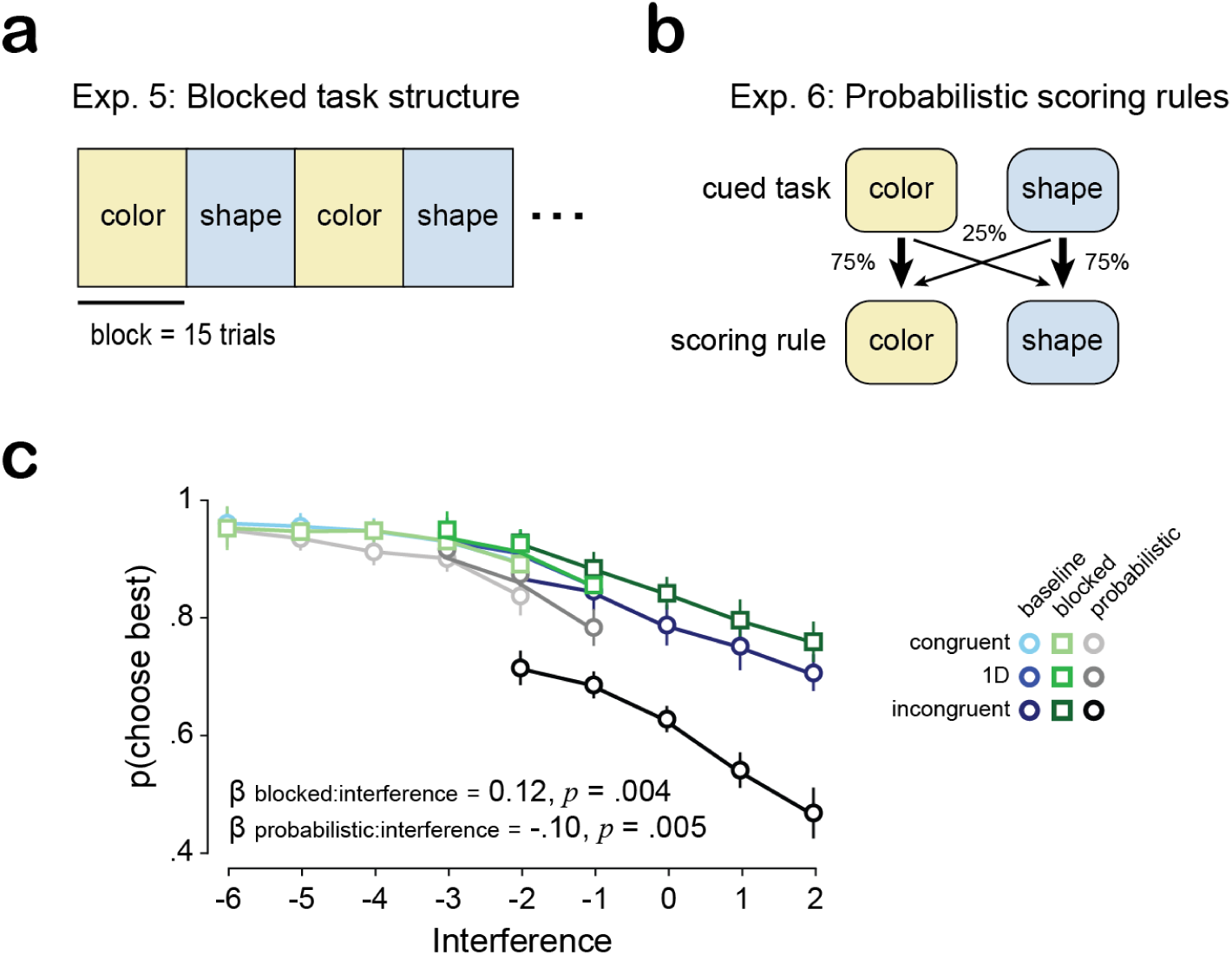
Experiments testing model predictions. **(a)** Using the original task design but running the experiment in blocks of color and shape trials. **(b)** Using the original task design with probabilistic scoring rules – meaning that only 75% of the time is the cued dimension the one choices are scored on. **(c)** Choice accuracy as a function of interference and experiment (blocked and probabilistic scoring). Note: markers and error bars denote the mean and bootstrapped 95% confidence intervals, respectively. The inset statistics are the results of a mixed effects logistic regression where *Task* was coded categorically with the baseline experiment used as the reference level.

In Experiment 6 (N = 50), we aimed to *increase* irrelevant feature weight by making that feature partially relevant to payoff. Using the original interleaved design, we applied a probabilistic scoring rule in which the cued feature determined the score on 75% of trials and the uncued feature determined it on the remaining 25% (**Fig. 5b**). For instance, on a trial cued as “color, ” the outcome would reflect the chosen option’s color 75% of the time and its shape the other 25%. We reasoned that because the irrelevant feature now determined payoff on a portion of trials, fully ignoring it was no longer optimal, and participants should place more weight on it. We therefore predicted that probabilistic scoring would increase irrelevant feature weight and amplify interference.

We tested these predictions by fitting a mixed effects logistic regression where choice accuracy varied as a function of interference and task (baseline [Exp. 1, **Fig. 1a**], blocked (**Fig. 5a**), and probabilistic scoring; **Fig. 5b**). *Task* was coded as a categorical variable and the baseline experiment served as the reference level, such that the coefficients for the blocked and probabilistic experiments reflect how each manipulation altered performance relative to baseline. Despite the blocked design affording participants a more predictable environment in which they could anticipate the upcoming relevant feature and pre-commit attentional weight accordingly, blocking did not significantly affect overall accuracy (β = 0.09, *p* = .43). In contrast, probabilistic scoring markedly reduced overall accuracy, particularly during incongruent trials (β = −1.25, *p* < .001).

Examining the task-interference interaction effects revealed that the two manipulations shifted the interference slope in opposite directions: blocking attenuated the effect of interference on choice accuracy (β = 0.10, *p* = .005) whereas probabilistic scoring amplified it (β = −0.13, *p* = .004). This is consistent with the idea that because the uncued feature now determined the payoff on a random quarter of trials, participants could no longer treat it as fully irrelevant and instead placed greater weight on it. Together, these results suggest that interference during feature-based valuation reflects a deep constraint on how people weigh relevant against irrelevant information, one that partially resists anticipatory suppression but is readily amplified when the reliability of the attentional state cue is reduced. This is consistent with precision-weighting accounts in which uncertainty about which information source is relevant leads to changes in how information is integrated across available features^38^.

### Experiment 7 – linking feature-based valuation to psychiatric symptoms

Difficulty ignoring irrelevant information is a hallmark of a number of psychiatric conditions ^24,39–42^. For example, people with addiction struggle to ignore drug-predictive cues despite conscious intention to abstain ^18–20,43^. Patients with obsessive compulsive disorder (OCD) struggle to ignore specific environmental features (e.g. asymmetrical objects, contaminated surfaces) that trigger compulsions, even while recognizing these stimuli as objectively irrelevant or harmless ^44,45^. The common thread across these conditions is a failure to filter irrelevant sensory information when constructing the value representations that guide behavior — precisely the computation our feature-based valuation model quantifies. Therefore, we next tested whether individual differences in susceptibility to irrelevant information during feature-based valuation correlated with psychiatric symptom severity.

We recruited an additional 106 human subjects to complete both our feature-based valuation task and a battery of clinically validated psychometric assessments of attention-deficit/hyperactivity disorder (ASRS) ^46^, obsessive-compulsive disorder (OCI-R) ^47^, depression (Zung SDS) ^48^, alcoholism (AUDIT) ^49^, and drug use (DAST) ^50^ severity. We fit our computational model to each participant to quantify the extent to which irrelevant information influenced their decisions (**Fig. 6a**). Because psychiatric disorders share overlapping symptoms and rarely map onto single, isolated diagnoses, we adopted a transdiagnostic, dimensional approach in line with the NIMH Research Domain Criteria (“RDoC”) framework ^51^. Therefore, rather than treating each assessment as an independent measure, we conducted factor analysis across all psychometric scale items to extract a smaller set of latent symptom dimensions that cut across traditional diagnostic boundaries. From analyzing the eigenvalues associated with different numbers of factors, we found that a three factor model provided the most parsimonious account of the data (**Fig. 6b,c**). Examining how the psychometric scale items loaded onto these factors revealed that they corresponded to psychiatric phenotypes associated with (1) OCD, (2) comorbid ADHD and depression, and (3) substance abuse (**Fig. 6c**). We then correlated each participant’s fitted irrelevant feature weight with their factor scores. We observed significant, positive correlations between irrelevant feature weight and factors related to OCD (*r* = .28, *p* = .005) and substance abuse (*r* = .32, *p* < .001) but not ADHD/depression, which trended negatively (*r* = -.16, *p* = .1) (**Fig. 6d**).

**Figure 6.**
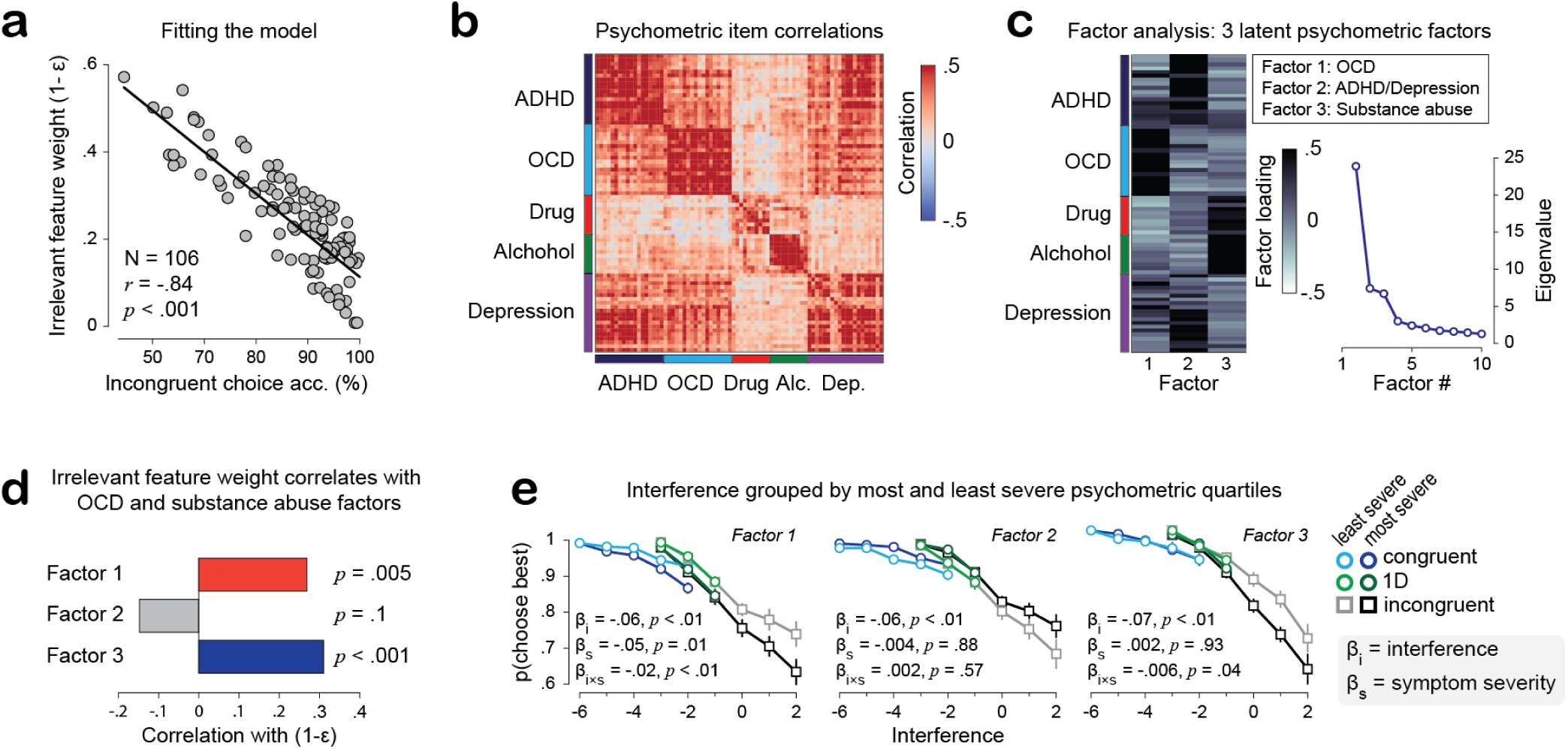
Correlating feature-based valuation with psychiatric symptom severity. **(a)** Fitting our computational model and extracting irrelevant feature weights for each of 106 new human participants. **(b)** Correlations between individual psychometric scale items. **(c)** Loadings for each psychometric scale item on each of 3 latent factors. The inset shows the eigenvalues for models with different numbers of factors. There was a clear elbow where models with more than 3 factors did not substantively improve fit. **(d)** Individual differences in irrelevant feature weight are positively correlated with psychiatric symptom severity related to obsessive compulsive disorder and substance abuse. **(e)** Choice accuracy as a function of interference and psychiatric symptom severity for each factor. Symptom severity was defined as the top and bottom quartiles of the factor scores. Markers and error bars denote means and bootstrapped 95% confidence intervals, respectively.

To visualize these findings, we plotted choice accuracy as a function of interference for participants in the top and bottom quartiles of each factor’s symptom-severity scores (**Fig. 6e**). Consistent with the correlations, participants high in OCD and substance-abuse symptoms showed steeper declines in accuracy with increasing interference than those low in these symptoms (i.e. significant negative severity × interference interactions in mixed-effects regression models). Taken together, these results suggest that compulsive and addictive symptoms may, in part, reflect an impaired ability to appropriately weigh relevant against irrelevant sensory information when assigning value to choice options.

## Discussion

We found that task-irrelevant sensory information interferes with how values are assigned to choice options. The sign of this influence depended on congruence: during congruent trials, where the relevant and irrelevant features favored the same option, interference was constructive, facilitating faster, more accurate choices. In contrast, during incongruent trials, where the two features conflicted, interference was destructive, leading to slower, less accurate choices.

This interference was not an artifact of task-switching, working memory, or semantic demands, but instead reflected a bottleneck in how people weigh relevant against irrelevant information during value-based choice. A generative model captured this susceptibility in a single parameter, the irrelevant feature weight. Critically, individual differences in this parameter tracked the severity of psychiatric symptom dimensions related to compulsivity and substance abuse. This suggests that these symptoms may, in part, reflect impairment in the ability to appropriately weigh relevant versus irrelevant sensory information when assigning value to choice options – a computation we term feature-based valuation.

The interference we observe is reminiscent of the classic Stroop task ^52,53^, in which participants name the ink color of a word while ignoring its meaning. However, feature-based valuation likely engages a different form of cognitive control. To illustrate, consider the Stroop stimulus “blue” printed in red ink: although “red” is the correct answer (because the font color is red), most people struggle not to report “blue” because reading is an automatic process that conflicts with color-naming. The Stroop task has proven clinically valuable precisely because this automatic interference maps onto real-world cognitive failures in a range of psychiatric conditions, most notably ADHD and schizophrenia ^54–56^. Yet Stroop interference is perceptual and linguistic: the irrelevant dimension carries semantic meaning, not reward value. No analogous paradigm has existed for the motivational domain, where interference arises not from automatically processed word meanings but from competing reward values. This distinction is not merely conceptual: Stroop performance does not predict OCD symptom severity ^57,58^ and predicts addiction severity only when the words themselves are drug- or alcohol-related ^31–33,59^ – suggesting these effects reflect cue reactivity rather than a general failure of distractor suppression ^59^. In contrast, feature-based valuation tracked symptom severity using affectively neutral stimuli, suggesting that it reflects a distinct, motivational form of interference and cognitive control.

One interpretation of our results is that the interference we observed reflects a competition between two mechanisms of attentional prioritization: top-down control ^9–11^ and automatic attentional capture via selection history ^14–17^. In our task, top-down control should direct valuation toward the cued feature, while the value of the uncued feature – which at other times was relevant and reward-predictive – exerts a competing pull. The irrelevant feature weights in our computational model may therefore index how effectively top-down control could override selection history biases. Notably, this competition appears dissociable from cognitive flexibility, *per se*: we observed clear interference effects both on trials where participants repeated the same relevant dimension and on trials where they switched between dimensions. This suggests that top-down control and selection history mechanisms compete for attentional control on each trial.

Our results also bear on an ongoing debate about how the brain represents value during multi-attribute choice. The traditional view holds that the brain computes an integrated value for each option by combining its attributes, then compares these integrated values to choose ^28,60,61^. However, recent work has challenged this view, suggesting that decisions instead proceed through attention-dependent comparisons of individual attributes rather than through comparison of integrated option values ^62,63^. Our findings are consistent with an attention-weighted account but indicate that attentional selection is incomplete: the irrelevant attribute was not excluded and contributed to it in proportion to its value.

This highlights a distinctive aspect of our task: unlike most multi-attribute choice tasks where each attribute (e.g. the amount, probability, or identity of reward) is relevant to determining an option’s overall value, the optimal policy in our task was to entirely ignore the uncued dimension. Our modeling revealed a spectrum of individual differences in how people deviate from this optimal policy. Some people performed near-optimal, assigning little-to-no weight to the irrelevant feature while others allowed the irrelevant dimension to exert considerable influence. Because weighting the irrelevant dimension was unambiguously suboptimal in our task, this variation reflects individual differences in susceptibility to irrelevant information when assigning value to choice options.

Interestingly, we found that individual differences in susceptibility to irrelevant information were positively correlated with psychiatric symptom severity related to OCD and substance abuse but not with a joint ADHD-depression factor. The emergence of a combined ADHD–depression factor is itself unsurprising as depression is among the most common comorbidities of adult ADHD ^64–66^. Given that distractibility is a defining feature of ADHD ^67,68^, we were initially surprised that this factor was not positively associated with impaired feature-based valuation (in fact, the correlation trended negatively). However, the distractibility characteristic of ADHD is more associated with difficulty maintaining focus and “staying on task” over extended periods – for example, trouble finishing projects, staying organized, and remembering appointments – as compared to the rapid shifts of attention required by our task. Indeed, the Adult ADHD Self-Report Scale ^46^, a widely used and clinically validated measure of adult ADHD symptoms ^46,69,70^, predominantly assesses difficulties in sustaining focus for longer durations. That we used this same, well-validated scale and found no significant relationship with ADHD symptom severity suggests that feature-based valuation engages a distinct, momentary form of attentional control different from the sustained-attention difficulties that characterize ADHD. That our results trended towards a negative correlation could suggest that the rapid shifts of attention required by our task may even have been facilitated by ADHD symptoms.

The specificity of our finding – that impaired feature-based valuation is associated with increased compulsive and substance abuse behaviors – complements a prior computational psychiatry study by Gillan and colleagues ^71^. In this study, the authors used reinforcement learning models to show that a transdiagnostic factor related to “compulsive behavior and intrusive thought ” is associated with reduced “goal-directed control “. Similarly, we found that a related symptom profile was associated with reduced control over feature-based valuation. Whereas Gillan and colleagues identified a failure to exert goal-directed control over action, we identify a failure to exert control over attentional prioritization and, subsequently, how values are assigned to choice options. Together, these studies illustrate how the computational psychiatry approach can identify clinically relevant components of cognitive control.

Our work is the first step towards bridging two literatures that have largely developed in parallel: feature-based attention and value-based choice. Neurophysiological research on feature-based attention has shown that attending to a stimulus feature enhances its neural representation while suppressing others ^35,72,73^. Research on value-based choice has focused on how the values of options are computed and compared to guide decisions, typically assuming that the relevant information has already been selected or otherwise integrated ^28,62,74^. Feature-based valuation sits at the intersection of these processes with the goal of understanding how attentional weighting of stimulus features shapes the values that ultimately drive choice. Our results show that this weighting is imperfect: that irrelevant features are not fully suppressed and intrude on valuation. Moreover, we show that individual differences in feature-based valuation are clinically meaningful, tracking the severity of compulsive and addictive psychiatric symptoms. There remains much to learn about how the brain selects which information to use during value-based choice and how this selection breaks down in disease.

## Methods

### Ethical approval

This study was reviewed and approved by the Institutional Review Board at the University of Texas at Austin. All participants provided informed consent before beginning the experiment. Participants were informed that their participation is voluntary and that they could withdraw at any time without penalty. No personally-identifiable information was collected.

### Participants

#### Recruitment

Across 7 experiments we recruited 406 participants through prolific.com, an online platform. We accepted any participant that resides in the United States, is fluent in English, has normal or corrected-to-normal vision, and is at least 18 years old. Experiments 1-6 took approximately 25 minutes to complete and participants were compensated $5 ($12/hour) for their time. Experiment 7, which included both our feature-based valuation task along with a battery of psychometric assessments, took approximately 45 minutes to complete and participants were compensated $9 ($12/hour) for their time.

#### Demographics

Table 1 below shows the demographics of participants who participated in each experiment.

#### Exclusion criteria

We excluded participants on the basis of task performance, engagement, and completion of the entire experiment. Our performance criterion was based on the “1D” trials of the feature-based valuation task - trials where the choice options varied only along the relevant dimension. Participants that scored less than 65% across these 1D trials were excluded from further analysis. We assessed engagement through a series of explicit attention checks throughout the psychometric assessment. These checks were items in the psychometric assessments that explicitly instructed participants to make a certain response (e.g. “For this item, select ‘a lot’”). We included at least two such attention check items in each psychometric assessment and excluded participants who failed to answer all of them correctly. Our final dataset reflects subjects who met these criteria.

#### Online behavioral testing

We programmed our experiment using the jsPsych JavaScript library ^75^ and hosted it online through cognition.run. Cognition.run is an online platform for hosting behavioral experiments programmed in jsPsych that can integrate with Prolific, the platform where we recruited our participants. Prior work ^76,77^ demonstrated that jsPsych maintains high timing accuracy when running experiments online and is widely and routinely used for reaction time studies. As detailed below, we included attention checks to ensure high quality data.

### Feature-based valuation task

#### Overview and mechanics

The feature-based valuation task measures how people prioritize one sensory feature over another when assigning values to choice options. On each trial, participants were explicitly cued whether to make a subsequent choice based on the values associated with either the colors or the shapes of the choice options. We defined value as the number of points a given feature yielded. We used four value levels per feature-dimension (i.e. four colors and four shapes). Following the cue (600 ms) and a brief delay (200 ms), a randomly selected pair of choice options was presented. All possible combinations of colors and shapes were presented across the trials with the only constraint that there was always a best option along the relevant sensory dimension. Participants indicated their responses with keypresses: they pressed the “f” key to select the left option and the “j” key to select the right option. Participants were allowed only one response per trial. Upon responding, a ring appeared around the chosen option for 200 ms to visually indicate to the participant that their choice was registered. Then both options and the selection ring were extinguished and participants were shown a 1000 ms feedback screen that indicated the number of points their choice yielded. The message shown was “You won X points!”, where “X” corresponds to the number of points their choice yielded.

#### Training blocks

Prior to starting the main experiment, participants completed two short training blocks of 12 trials each designed to familiarize the participants with the features and their values. During the “color” training block, participants exclusively chose between pairs of octagons that varied only in their color; during the “shape” training block, participants exclusively chose between pairs of gray shapes (i.e. no color information). The octagons used for the color block and gray color used for the shape block were not used in the main experiment. In other words, only the learning about the specific feature values carried over to the main experiment. The order of the training blocks was counterbalanced across participants.

After the training blocks, participants completed a brief comprehension test about the mechanics of the experiment (what do pressing the “f” and “j” keys do? How should they choose when initially shown the “color” or “shape” context cue). Participants could only continue to the main experiment when they passed this comprehension test.

#### Experimental blocks

Participants then performed two blocks of 192 trials (384 total) where all combinations of colored-shapes were presented, with the constraint that there was always a best option along the relevant (cued) dimension. After the first block, participants took a self-paced break, and then completed the second block.

#### General statistics

Unless otherwise noted, all markers and error bars in the figures reflect the mean and bootstrapped 95% confidence intervals. We made substantial use of mixed effects logistic and linear regressions, which were carried out in python using the `pymer` ^78^ and `statsmodels` ^79^ packages. We verified that all regression models did not suffer from multi-colinearity by computing the variance inflation factor for each term in each candidate model. We only proceeded with models where all factors had variance inflation factors of approximately 1, indicating that there was minimal collinearity between model terms.

#### Interference index

To capture how relevant and irrelevant feature values jointly contributed to choice, we constructed an interference index from the value differences along each dimension. For each trial, we first identified the option favored by the relevant (cued) feature – i.e., the option with the higher relevant-feature value, which by task design was always uniquely defined. We then computed:

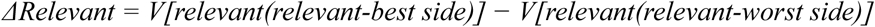

This quantity was always greater than zero since our task design ensured there was always a better option along the relevant dimension.

We then computed the irrelevant-feature value difference, signed relative to the same reference side:

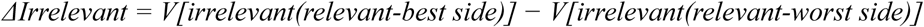

*ΔIrrelevant w*as positive when the irrelevant feature also favored the relevant-best side (congruent trials), negative when it favored the opposing side (incongruent trials), and zero when the irrelevant feature did not differ between the two options (1D trials).

We then defined the interference index as:

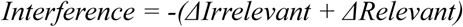

This index is negative on congruent trials, where the irrelevant feature reinforces the relevant feature, and increases on incongruent trials, where the irrelevant feature conflicts with the relevant feature. In other words, *Interference* increases as the feature-dimensions progressively favor different options such that lower values reflect agreement and higher values reflect conflict between them.

#### Generative model of feature-based valuation

We constructed a generative model of choice designed to capture the constructive and destructive interference effects observed in our behavioral data – namely, that irrelevant feature values facilitate choice when they agree with the relevant feature and impair choice when they conflict with it. The model formalizes this idea by treating the decision variable on each trial as a weighted sum of noisy relevant and irrelevant feature values, such that the irrelevant feature’s value is never fully excluded from the decision but instead exerts an influence proportional to its own value and to how strongly it is weighted. The model has two free parameters: a feature weight (ε), which governs the relative contribution of the cued (relevant) versus uncued (irrelevant) feature to the decision, and value noise (σ), which governs the stochasticity of the internal representation of each feature’s value.

On each trial, the model first samples a noisy internal estimate of each option’s feature values by drawing from a normal distribution centered on the true value:

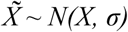

where *X* is the true (objective) value of a given feature and *σ* is the standard deviation of the noise added to its internal representation. This was applied independently to the relevant and irrelevant feature value of each option. The model then computed a decision variable (*DV*) as a weighted sum of the noisy relevant and irrelevant value differences between the two options:

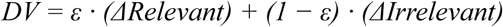

where *ΔRelevant* and *ΔIrrelevant* are the noisy value differences (best minus worst option, signed according to the relevant dimension) along the relevant and irrelevant feature dimensions, respectively, and ε is bounded between 0 and 1. ε = 1 corresponds to the optimal strategy, in which the irrelevant feature is given no weight and choice is determined entirely by the relevant dimension; ε = 0.5 corresponds to equal weighting of relevant and irrelevant information. We refer to the complementary quantity, 1 − ε, as the irrelevant feature weight, since it directly quantifies the degree to which an individual’s choices were influenced by task-irrelevant information.

This decision variable was then converted into a choice probability. Because each feature value is represented with Gaussian noise, the DV is itself a normally distributed random variable with mean:

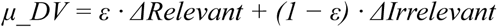

and standard deviation:

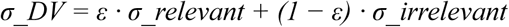

where *σ_relevant* and *σ_irrelevant* denote the standard deviation of the noisy value differences along each dimension (each equal to σ√2, reflecting the difference of two independently noisy value estimates). The probability of choosing the best option was then computed as the probability that the *DV* exceeds a decision criterion of zero:

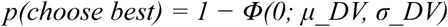

where *Φ* denotes the cumulative normal distribution function. In other words, because the *DV* is a distribution, the choice probability reflects the proportion of that distribution that exceeds the criterion of 0.

We fit the model separately to each participant by finding the parameter values (ε, σ) that maximized the likelihood of each participant’s observed trial-by-trial choices. Fitting was performed via maximum likelihood estimation using `scipy.optimize` in Python.

#### Psychometric scales and factor analysis

In Experiment 7, participants completed both our feature-based valuation task and a battery of clinically validated psychometric assessments of attention-deficit/ hyperactivity disorder (ASRS) ^46^, obsessive-compulsive disorder (OCI-R) ^47^, depression (Zung SDS) ^48^, alcoholism (AUDIT) ^49^, and drug use (DAST) ^50^ severity. Because psychiatric disorders share overlapping symptoms and rarely map onto single, isolated diagnoses, we adopted a transdiagnostic, dimensional approach in line with the NIMH Research Domain Criteria (RDoC) framework ^51^. Therefore,rather than treating each assessment as an independent measure, we conducted factor analysis across all psychometric scale items to extract a smaller set of latent symptom dimensions.

We based our factor analysis on the one established by Gillan et al. ^71^. First, we z-scored each individual survey item. We then employed factor analysis using the `FactorAnalyzer` python module, with an oblique rotation. Factor selection was based on Cattell’s criterion ^80,81^, where we identified a transition from horizontal to vertical (“elbow”) in the eigenvalues associated with each factor (see Scree plot inset in Fig. 6c) – indicating that there is little benefit to retaining additional factors. Our analysis indicated the existence of a 3-factor latent structure that we labeled “Compulsive behavior”, ADHD/Depression”, and “Substance Abuse” based on the pattern of individual item factor loadings (Fig. 6c).

## Code availability

All code supporting this project are publicly available on our lab’s github, which can be accessed at: https://github.com/elston-lab/human-feature-based-valuation

This repository contains the Javascript and stimuli to run the experiment as well as Jupyter notebooks for the analysis. Explicit instructions for how to reproduce the experiment are provided.

## Data availability

The data that support the findings of this study are available on our lab’s github, which can be accessed at: https://github.com/elston-lab/human-feature-based-valuation

## Acknowledgements

TWE is supported by a startup grant from the UT Austin College of Natural Sciences. MGM is supported by a T32 training grant from the National Eye Institute (T32EY021462). RKR is supported by a T32 training grant from the National Institute on Drug Abuse (T32DA018926). We thank Drs. Clara Starkweather, Preeya Khanna, Jarrod Lewis-Peacock, Ian Mackenzie, Alyssa Sanchez, Eric Hu, and Robbe Goris for helpful discussion and feedback on the manuscript.

